# Genomic and Chemical Diversity in *Cannabis*

**DOI:** 10.1101/034314

**Authors:** Ryan C. Lynch, Daniela Vergara, Silas Tittes, Kristin White, C.J. Schwartz, Matthew J. Gibbs, Travis C. Ruthenburg, Kymron deCesare, Donald P. Land, Nolan C. Kane

## Abstract

Plants of the *Cannabis* genus are the only producers of phytocannabinoids, terpenoid compounds that strongly interact with evolutionarily ancient endocannabinoid receptors shared by most bilaterian taxa. For millennia, the plant has been cultivated for these compounds, but also for food, rope, paper, and clothing. Today, specialized varieties yielding high-quality textile fibers, nutritional seed oil or high cannabinoid content are cultivated across the globe. However, the genetic identities and histories of these diverse populations remain largely obscured. We analyzed the nuclear genomic diversity among 340 *Cannabis* varieties, including fiber and seed oil hemp, high cannabinoid drug-types and feral populations. These analyses demonstrate the existence of at least three major groups of diversity, with European hemp varieties more closely related to narrow leaflet drug-types (NLDT) than to broad leaflet drug-types (BLDT). The BLDT group appears to encompass less diversity than the NLDT, which reflects the larger geographic range of NLDTs, and suggests a more recent origin of domestication of the BLDTs. As well as being genetically distinct, hemp, NLDT and BLDT genetic groups each produce unique cannabinoid and terpenoid content profiles. This combined analysis of population genomic and trait variation informs our understanding of the potential uses of different genetic variants for medicine and agriculture, providing valuable insights and tools for a rapidly emerging, valuable legal industry.

## Significance Statement

Despite millennia of cultivation and current widespread use across the globe, *Cannabis* is the only multi-billion dollar crop for which the genetic identities and origins of most varieties are unknown. As legalized cultivation of hemp and high-cannabinoid types continues to grow rapidly in the US and other countries, the need for a better understanding of the diversity and evolution of the species has increased. Through analyzing the genomes of 340 hemp, drug and feral *Cannabis* individuals, we found significant evidence for at least three major genetic groups. Importantly, each group produces distinct phytochemical profiles. Our results improve the understanding of genetically and chemically diverse *Cannabis* strains currently cultivated, and provide a roadmap for developing improved varieties.

## Introduction

Plants of the genus *Cannabis* (Cannabaceae; hemp, drug-type) have been used for thousands of years for fiber, nutritional seed oil and medicinal or psychoactive effects. Archaeological evidence for hemp fiber textile production in China dates to at least as early as 6,000 years ago (1), but possibly as early as 12,000 years ago (2), suggesting *Cannabis* was one of the first domesticated fiber plants. Archeological evidence for medicinal or shamanistic use of *Cannabis* has been found at Indian, central-Asian and middle-eastern sites (3), further illustrating the widespread extent of *Cannabis* utilization throughout human history. A central Asian site of domestication is often cited (4), although genetic analyses suggest two independent domestication events may have occurred separately (5).

*Cannabis* plants are usually annual wind-pollinated dioecious herbs, though individuals may live more than a year in subtropical climates (6) and monoecious populations exist (7). The taxonomic composition of the genus remains unresolved, with two species (*C*. *indica* and *C. sativa*) commonly cited (8), although *C. ruderalis* is sometimes proposed as a third species that contains northern short-day or auto-flowering plants (9). Monospecific treatment of the genus as *Cannabis sativa* L. is also common (10) and various alternative nomenclature schemes (e.g. *Cannabis sativa* subsp. *indica* var. *kafiristanica*) are sometimes referenced (4). Even though an extensive monograph on the genus has recently been published (11), limited genetic and experimental data leaves the questions of taxonomy unresolved (12, 13).

The geographical and ecological range of *Cannabis* is unusually broad, with cultivated populations growing outdoors on every continent except Antarctica in a wide range of environments from sub-arctic to temperate to tropical, and from sea level to over 3,000 meters elevation (14, 15). Feral or wild populations are also found as far north as the edge of the Arctic Circle in Eurasia, but are most common in well drained soils of temperate continental ecosystems in Eurasia and North America, while tropical populations are absent or rare (14). Perhaps unsurprising, given this diversity of habitats, the species contains extensive phytochemical diversity, particularly in cannabinoid and terpenoid profiles (5, 16), and also shows extensive diversity of morphological and life-history characteristics, further fueling debate regarding the taxonomic status and origins of *Cannabis* domestication.

One distinctive feature of the *Cannabis* genus is the production of a tremendous diversity of compounds called *cannabinoids*, so named because they are not produced at high levels in any other plant species (17). Cannabinoids are a group of at least 74 known C21 terpenophenolic compounds (18, 19) responsible for many reported medicinal and psychoactive effects of *Cannabis* consumption (20). Some estimates for the total number of phytocannabinoids range to well over a hundred (21), though this number includes breakdown products as well as compounds found at extremely low levels. The plants produce a non-psychoactive carboxylic acid form of these compounds, with heating required to convert cannabinoids into the psychoactive decarboxylated forms. Interestingly, these compounds have pronounced neurological effects on a wide range of vertebrate and invertebrate taxa, suggesting an ancient origin of the endocannabinoid receptors, perhaps as old as the last common ancestor of all extant bilaterians, over 500 MYA (22). The plant compounds thus produced have the potential to affect a broad range of metazoans, though their ecological functions in nature are not well understood. Indeed, suggested roles for these compounds include many biotic and abiotic defenses, such as suppression of pathogens and herbivores, protection from UV radiation damage, and attraction of seed dispersers. These hypotheses about the selective benefits of cannabinoid production remain speculative, as none have been conclusively verified to date. We do know more, however, about the more recent evolution of the plants under human cultivation.

High delta-9-tetrahydrocannabinolic acid (THCA) (23) content has been selected for in many strains due to its potential to be converted to delta-9-tetrahydrocannabinol (THC), which has potent psychoactive (24), appetite-stimulating (25), analgesic (26) and antiemetic (27) effects. These effects are mediated through interactions with human endocannabinoid CB1 receptors found in the brain (28), and CB2 receptors, which are concentrated in peripheral tissues (29). Other THC receptor binding locations are hypothesized as well (30). After several decades of accelerated clandestine cultivation technique and breeding improvements, some modern strains can now yield dried un-pollenated pistillate inflorescence material that contains over 30% THCA by dry-weight (31). However, other cannabinoids may also be present in high concentrations. In particular, high cannabidiolic acid (CBDA) plants were historically used in some hashish preparations(32) and are presently in high demand as an anti-seizure therapy (33). In contrast with THC, which acts as a partial agonist of the CB1 and CB2 receptors, CBD does not have as strong psychoactive properties, and instead has antagonist activity on agonists of the CB1- and CB2-receptors (34). Thus, the two most abundant cannabinoids produced in *Cannabis* have, to some degree, opposing neurological effects.

THCA and CBDA are alternative products of a shared precursor, CBGA (35). A single locus with co-dominant alleles was proposed to explain patterns of inheritance for THCA to CBDA ratios (7, 36). However more recent quantitative trait loci (QTL) mapping experiments (37), expression studies (38) and genomic analyses (10) paint a more complex scenario with several linked paralogs responsible for the various THCA and CBDA phenotypes. Other cannabinoids such as cannabigerol (CBG) (39), cannabichromene (CBC) (40) and delta-9-tetrahydocannabivarin (THCV) (41) demonstrate pharmacological promise, and can also be produced at high levels by the plan t(42–44). Additionally, *Cannabis* secondary metabolites such as terpenoids and flavonoids likely contribute to therapeutic or psychoactive effects (2), such as β-myrcene, humulene and linalool proposed to produce sedative effects associated with specific strains (45).

In this study, plants that produce low levels of total cannabinoids are herein referred to as hemp, while high cannabinoid producing varietals are described as drug-type strains. Legal definitions often use a maximum THCA threshold to delineate hemp from drug-types, thus some high CBDA producing strains are categorized as hemp. However this definition ignores the broader traditional usage of hemp for fibers or seed oils and historical presence of CBDA-producing alleles in some drug-type populations (32). Additionally, hemp strains have a distinct set of growth characteristics (46), with fiber varieties reaching up to 6 meters in height during a growing season, exhibiting reduced flower set, increased internodal spacing and lower total cannabinoid concentration per unit mass compared to drug-type relative. Despite the widespread prohibition of drug-type *Cannabis* cultivation from the 1930s to present (47), hemp cultivation and breeding continued in parts of Europe and China though this period, and experienced a brief comeback during World War II in the USA through the Hemp for Victory campaign. Studies to date have found hemp varieties are genetically distinct from drug-type strains (10), though interestingly Hillig (5) found broad leaflet southeastern Asian hemp landraces to be more closely related to Asian drug-type strains than to European hemp strains.

*Cannabis* has a diploid genome (2n = 20), and an XY/XX chromosomal sex-determining system(48). The genome size is estimated to be 818 Mb for female plants and 843 Mb for male plants (49). Currently, a draft genome consisting of 60,029 scaffolds is available for the Purple Kush (PK) drug-type strain from the National Center for Biotechnology Information. Additional whole genome data is available from NCBI for the Finola and USO31 hemp strains. Various reduced representation genome, gene and RNA sequence data are also available from NCBI. Presently *Cannabis* is the only multi-billion dollar legal crop without a sequence-based genetic linkage or physical genome map. Indeed, the first genetic map for the species, using AFLP and microsatellite markers, was only recently published, providing for the first time, quantitative trait mapping of cannabinoid content and other traits (37).

Initial studies of *Cannabis* genetic diversity examined either many samples with few molecular markers (5) or whole genome wide data for relatively few samples types (10). Sawler et al. (50) recently published a survey of *Cannabis* genomic diversity, using a reduced genomic representation strategy to evaluate 81 marijuana (drug-type) and 43 hemp strains. The aim of this present study is to assess the genomic diversity and phylogenetic relationships among 340 total *Cannabis* plants that have distinct phenotypes, and that were described *a priori* by plant breeders as various landraces, *indica, sativa*, hemp and drug-types, as well as commercially available hemp and drug-types with unclear pedigrees. We have combined data from existing sources and generated new data to create the largest sample set of *Cannabis* genomic sequence data published to date. These data and analyses will continue to facilitate the development of modernized breeding and quality assurance tools, which are lacking in the nascent legal *Cannabis* industry.

## Results and Discussion

**Sequencing and SNPs**. Summary information and raw sequencing libraries are publically available from the NCBI short read archive (accessions pending). Detailed information about all samples can be found in Dataset S1 and examples of wide and narrow leaflet forms are shown in Figure 1. Of the 466,427,059 non-ambiguous base pairs in the PK reference, 77,810,563 bps were removed due to excess self-similarity (≥ 97 % identity and ≥ 500 bps length, Figure S1). After this filter, the total single copy portion the PK reference within the combined coverage levels for all 67 WGS samples of 326x – 401x, a 95% Poisson confidence interval around a 362x mean, was 71,236,365 bps (Figure S1). After quality (Q), genotype quality (GQ), allele frequency (AF), missing data, biallelic and ambiguous base filters, the following SNP counts remained: 491,341 WGS, 2,894 GBS (this study), SNPs 4,105 GBS Sawler (50). Forty-five SNPs overlapped both GBS datasets, and the WGS samples.

**Figure 1.**
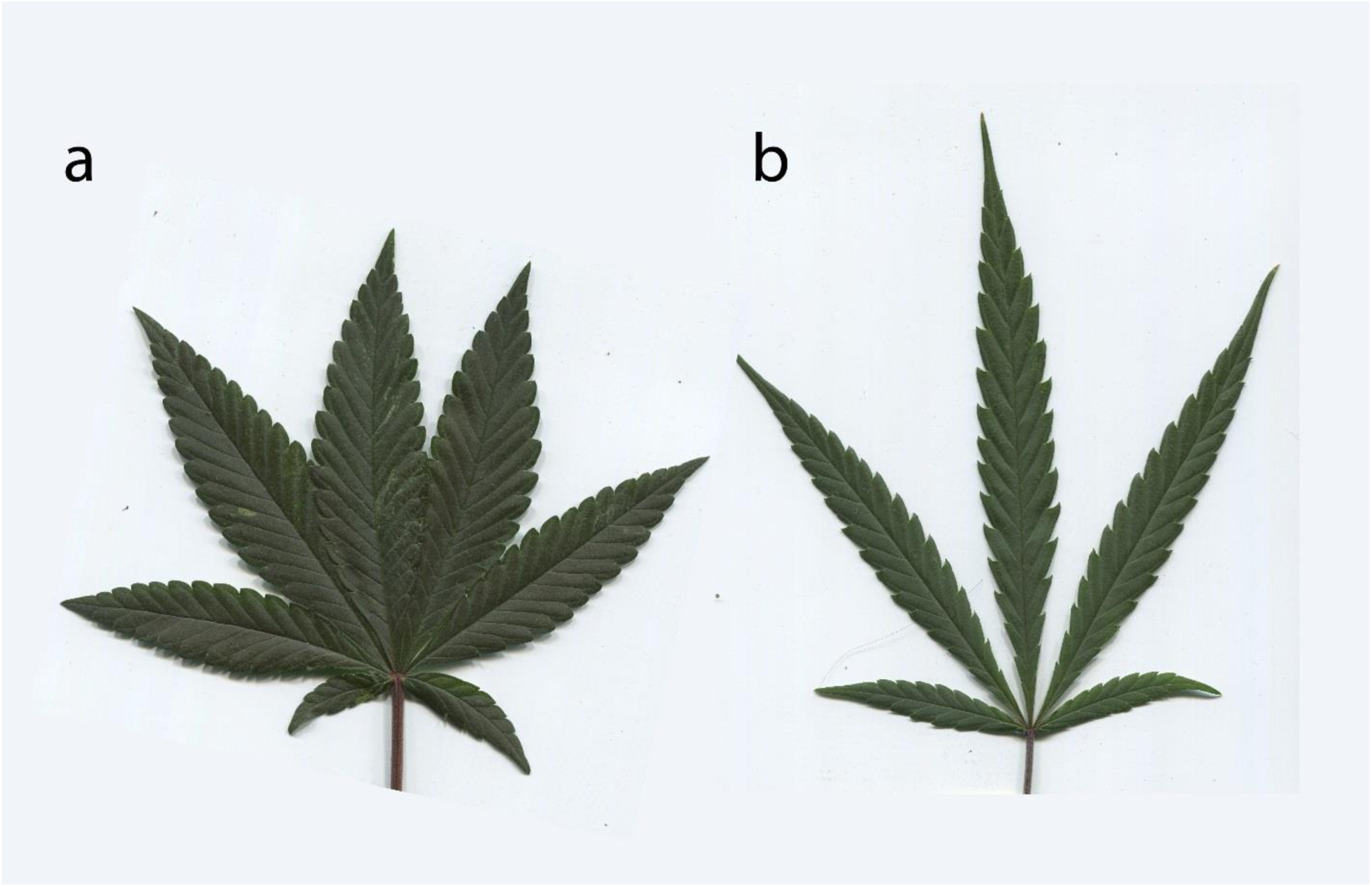
Example of broad leaflet type (a, R4) and narrow leaflet type (b, Super Lemon Haze) strains. Photograph credits: D. Vergara.

**Phylogenetic Relationships**. Bifurcating trees are commonly used to model mutation driven divergence and speciation events. Whole genome wide sequence datasets include information about recombination, hybridization, and gene loss or genesis events, some of which may be incongruent with one and other (51). Phylogenetic networks can represent incompatible phylogenetic signals across large character matrices in a visually informative manner. Figure 2 contains 195 *Cannabis* samples including WGS and GBS data, and shows that all European hemp strains form a distinct clade, separated from drug-type strains by a consistent band of parallel branches. Broad leaflet drug-type strains clustered with purported Afghan Kush landrace samples (Dataset S1 and Figure S3), while narrow leaflet drug-type strains appear to contain several groups with only faint visible distinctions between them, perhaps influenced by the inclusion of hybrid strains in the analysis.

**Figure 2.**
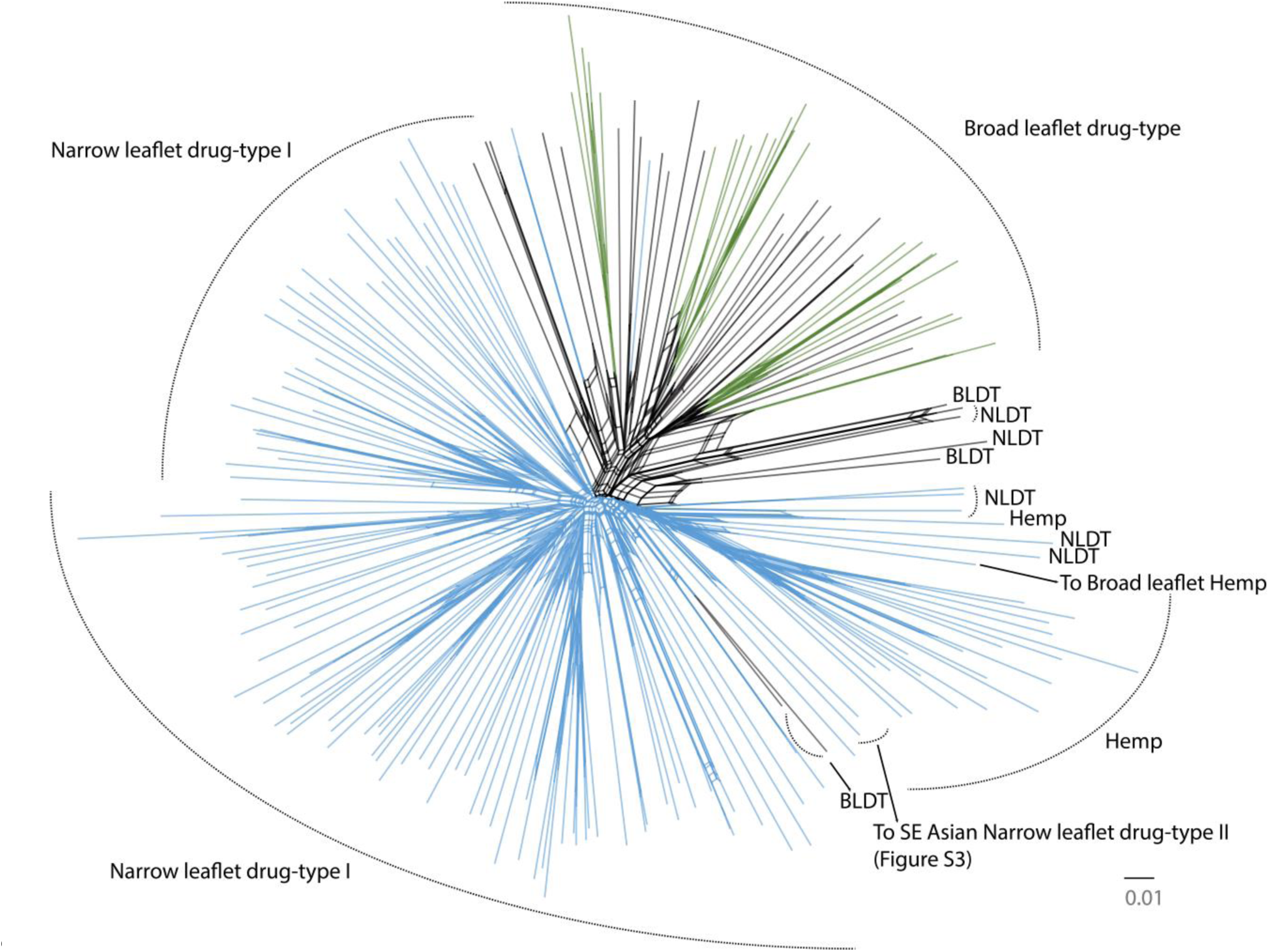
Phylogenetic neighbor network of a 2,894 SNP alignment from the single-copy portion of the *Cannabis* genome. Clade names on the periphery were inferred via FLOCK (where K ≥ 3 was most likely). Colored branches indicate fastStructure population membership of ≥ 70% assignment (where K=2 was most likely). NLDT = Narrow Leaflet Drug-Type and BLDT = Broad Leaflet Drug-Type. To SE Asian NLDT II points to Dr. Grinspoon and Somali Taxi Cab samples. To Broad Leaflet Hemp points to a Chinese hemp sample. A high-resolution version of this figure that includes each sample name is available from: https://figshare.com/articles/Cannabis_Tree/1585470/4

We found significantly more heterozygosity in drug-type strains than in hemp varieties (31 *%* v 22 %, p < 0.001, two-tailed Mann-Whitney U-test, Table 1). This likely reflects the widespread hybridization of strains in North America during the transition to indoor cultivation of drug-types starting in the 1970s (52), as well as the extensive reliance on clonal propagation for indoor commercial cultivation, which does not require trait stable seed stock. Conversely, fiber and seed oil hemp are grown on multi-acre scales that have necessitated the stabilization of agronomically important traits in seed stocks, likely leading to reduced heterozygosity at some loci.

## Group Genetic Information

**Table 1.** Summary of genetic distance, heterozygosity and F_st_ information for major *Cannabis* groups. * = significantly different (p < 0.001, two-tailed Mann-Whitney U-test).

|  | Mean Within Distances | Heterozygosity % |
| --- | --- | --- |
| Hemp | 0.195 | 0.22* |
| All Drug-types | 0.244 | 0.31* |
| NLDT | 0.237 | 0.32 |
| BLDT | 0.221 | 0.30 |
|  | Mean Between Distances | F<sub>ST</sub> |
| Hemp v. All Drug-types | 0.273 | 0.098530 |
| Hemp v. NLDT | 0.269 | 0.091679 |
| Hemp v. BLDT | 0.281 | 0.10131 |
| NLDT v. BLDT | 0.258 | 0.036156 |

**Population Structure**. To determine the statistical likelihood of various population scenarios represented in our samples, we first applied the FLOCK model to our data set of 195 GBS and WGS *Cannabis* samples, which is an iterative reallocation clustering algorithm that does not require non-admixed individuals to make population assignments (53). Using the K-partitioning method suggested by the authors (53), we determined that K ≥ 3, after testing K values of one to eight (Table 2 and peripheral population names in Figure 2). FLOCK was able to assign all samples to one of the three populations, although it does not calculate admixture proportions. Sample population assignments were largely consistent with the known history of these samples, and appear visually consistent with MDS analysis (Figure S2). For example all fiber and seed oil hemps were assigned to an exclusive population, with the exception of sample AC/DC, a high CBDA producing variety, with likely hybrid hemp origins (Figure 2, Table 2).

**Table 2.** Sample names and FLOCK assignment to three groups, represented with different cell colors. Green are BLDT, blue are NLDT and yellow are hemp.

|  |  |  |  |  |
| --- | --- | --- | --- | --- |
| A-train | Original_Sour_Diesel | C36 | H11 | Schemp |
| Afghan_Kush | Phantom_Cookies | C37 | H5 | Skunk_#1 |
| Afghan_Kush | Platinum_OG | Canna_Tsu | Harlequin | Somali_Taxi_Cab |
| Afghan_Kush | Purple_Kryptonite | Cannatonic | Hash_Plant | Spectrum-11 |
| Afghan_Kush | Purple_Kush | CBD_Diesel | Hawaiian | Spectrum-14 |
| Afghan_Kush | Purple_Urkle | CBD_Shark_F-6 | Holy_Grail | Super_Lemon_Haze |
| Boss_Hogg | R4 | CBD-0 | Holy_Grail_Kush | Sweet_Afghani_Delicious |
| Bubba_Kush | San_Fran_Valley_OG_Kush | CBD18 | Jack_47 | Sweet_Skunk |
| Char_Tango | Screaming_Haze | Charlottes_Web | Jack_Flash | Tangerine_Haze |
| Chem_91' | SFV | Cheese_Quake | Jack_Herer | Train_Wreck |
| Chem91 | Skywalker_OG | Cherry | Jack_Herrer | Trainwreck |
| Chemdawg | Snowcap | Cherry_Afghan | Jack_Herrer | Violator_Kush |
| Chocolate_Kush | Snowcap | Chocolope | Jack_Skellington | White_Cookies |
| Crippd_Out_Cookies | Sour_Diesel | Chocolope | Juicy_Fruit | White_Widdow |
| Cript_out_Cookies | Sour_Patch_Kush | Colombia_Rio_Negro | Lebanese | White_Widow |
| Dead_Head_OG | Sour_Willie | Critical_Kush | Lemon_Skunk | Wonder_Woman |
| Dog_Walker | Sshrek | Critical_Kush | Liberty_Haze | XJ_13 |
| Flo | The_Sauce | Critical_Mass | Lions_Tabernacle | AC/DC |
| Girl_Scout_Cookie_#6 | The_Sauce | Dr_Grinspoon | Low_Ryder | AZ_Star_#1 |
| Girl_Scout_Cookies | Tora_Bora | Durban_Poison | LSD | Carmagnola |
| Girl_Scout_Cookies | WAF_B | Durban_Poison | Mad_Cow | Carmagnola |
| Goast_Train_Haze | 4-Jack | Easy_Sativa | Mango_Stomper | Carmagnola |
| Grapefruit | Afghan_Mango | Exodus_Cheese | Maui_Waui | Carmagnola |
| Guido_OG | Afghan_Mango | G13 | Mazar | Carmagnola |
| Headband | Afghan_Mango | G13_Haze-31 | Medical_Mass | Carmagnola |
| Hindu_Kush | Alaskan_Thunderfuck | Gin | Melon_Gum | Chinese_hemp |
| Kandy_Kush | Appalachian_Mad_Sun | Girl_Scout_Cookies | Mexican_E | Dagestani_hemp |
| King_Chem | Auto_AK47 | Glass_Slipper | MO | EuroOil_2 |
| King_Louie_Cookies | B-5 | Golden_Goat | Nuclear_Fruit | Feral_Kansas |
| Kool_Aid_Kush | BC_HQ | Grand_Daddy_Purps | Otto | Feral_Nebraska |
| Kosher_Kush | Black_Cherry | Grape_AK-47 | Peaches_and_Cream | Feral_Nebraska |
| Kosher_Kush | Black_Jack | Grape_Ape | Pineapple | Finola |
| Kosher_Kush_#1 | Blue_Cheese | Grape_Ape | Pineapple_Express | J7 |
| Kunduz | Blue_Dream | Grape_Kush | Pink_Lady | J20 |
| Larry_OG | Blue_Dream | Grape_Kush | Pre-98_Bubba_Kush | J28 |
| Medibud | Blue_OG | Green_Crack | Purps | Kompolti_1 |
| OG_18 | Blueberry_DJ | Green_Crack | R4 | Kompolti_2 |
| OG_Kush_1 | Bubble_Gum | Green_Mandarine | Red_Purps | Sievers_Infinity |
| Old_Skool_OG | Bubble_Gum_XL | Green_Poison | Rocky_Mountain_Blueberry | US031 |

Additionally we applied the admixture model based Bayesian clustering method of fastStructure to the same 195 samples (54). The most likely population structure analysis of K=2 (Figure 2, Dataset S1), shows consistent separation between BLDT and NLDT and hemp strains. Some hemp and NLDT strains were each assigned with near 100% population membership to the same population (Figure 2, light blue samples, Dataset S1), despite the clear separation visualized in the tree and statistically significant mean between-group genetic distance measured (Table 1). The separation of BLDT and NLDT strains into fastStructure populations was stable when hemp samples were excluded from the analysis (Dataset S1). Sawler et al. (50) used fastStructure to delineate hemp from drug-types as the major division of *Cannabis* diversity, and found two drug-type sub-groups within their samples when hemp types were excluded from the analysis. Likewise using a smaller dataset, Lynch (55) found support for K=3, consisting of two separate drug-type populations and hemp types, using the original Structure implementation (56) and the Evanno method to select the best value of K (57). However, we caution that despite many claims for the availability of “landrace genetics” (strains) from *Cannabis* producers, breeders and seed sellers, these may or may not represent non-admixed individuals (52)—a situation that can be problematic for the Structure and fastStructure approaches (56).

The GBS samples from Sawler et al. (50) appear to contain an additional divergent NLDT clade, with likely SE Asian origins (Supplementary Figures 3 and 4), that did not emerge from our main analyses. Due to very limited overlap between sequence fragments from the two GBS datasets, which results from using different restriction enzymes, we were required to reanalyze the Sawler data in combination with only our 67 WGS samples. A connection was made across the two GBS analyses to this SE Asian NLDT group through two WGS samples (Dr. Grinspoon and Somali Taxi Cab, Figure 2, Supplementary Figures 3) that were included in both sets of GBS analyses. Although only 45 SNPs overlapped between both types of GBS data and the WGS data, a phylogeny of this limited alignment also supports the existence of an additional distinct SE Asian NLDT clade (Figure S4). Collectively these analyses lend support to a total lower bound of four *Cannabis* populations, although clearly more extensive sampling with consistent sequencing is required to fully access standing biogeographic diversity.

**Tests of Tree Models**. To test hypotheses of tree-like evolution for the three genetic groups, we first applied the three-population test for admixture (58), and found no evidence for admixture in any of the pairwise comparisons (positive *f* statistic values). Next we constructed maximum likelihood trees based on the aggregate SNP frequencies for the three genetic groups and simulated a variety of ‘migration’ events (0-10), but no simulation produced non-zero migration graph edges (Figure 3). F_ST_ analysis shows little divergence among lineages for most loci, but a substantial number of highly-divergent regions are unique to each clade (Figure 4). This reinforces the importance of using many, high-quality, single-copy regions of the genome, rather than smaller numbers of loci that could lead to less resolution or even misleading results. Although lore (52), Figure 2 and Figure S2 strongly suggest at least some individuals have hybrid origins, these tree models for the overall SNP frequencies of the population groups inferred by FLOCK (Table 2) imply each group contains strong genetic signals from ancestral biogeographic gene pools.

**Figure 3.**
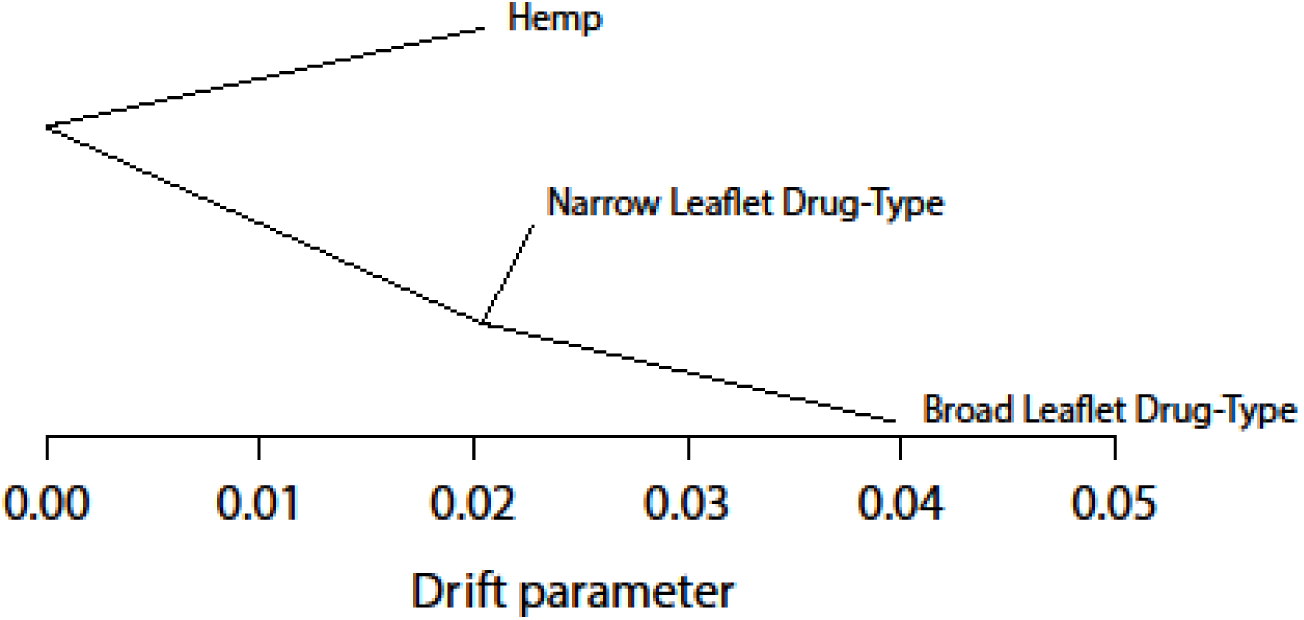
Maximum likelihood tree of three *Cannabis* populations. We found no evidence for extensive admixture or deviations from this tree model.

**Figure 4.**
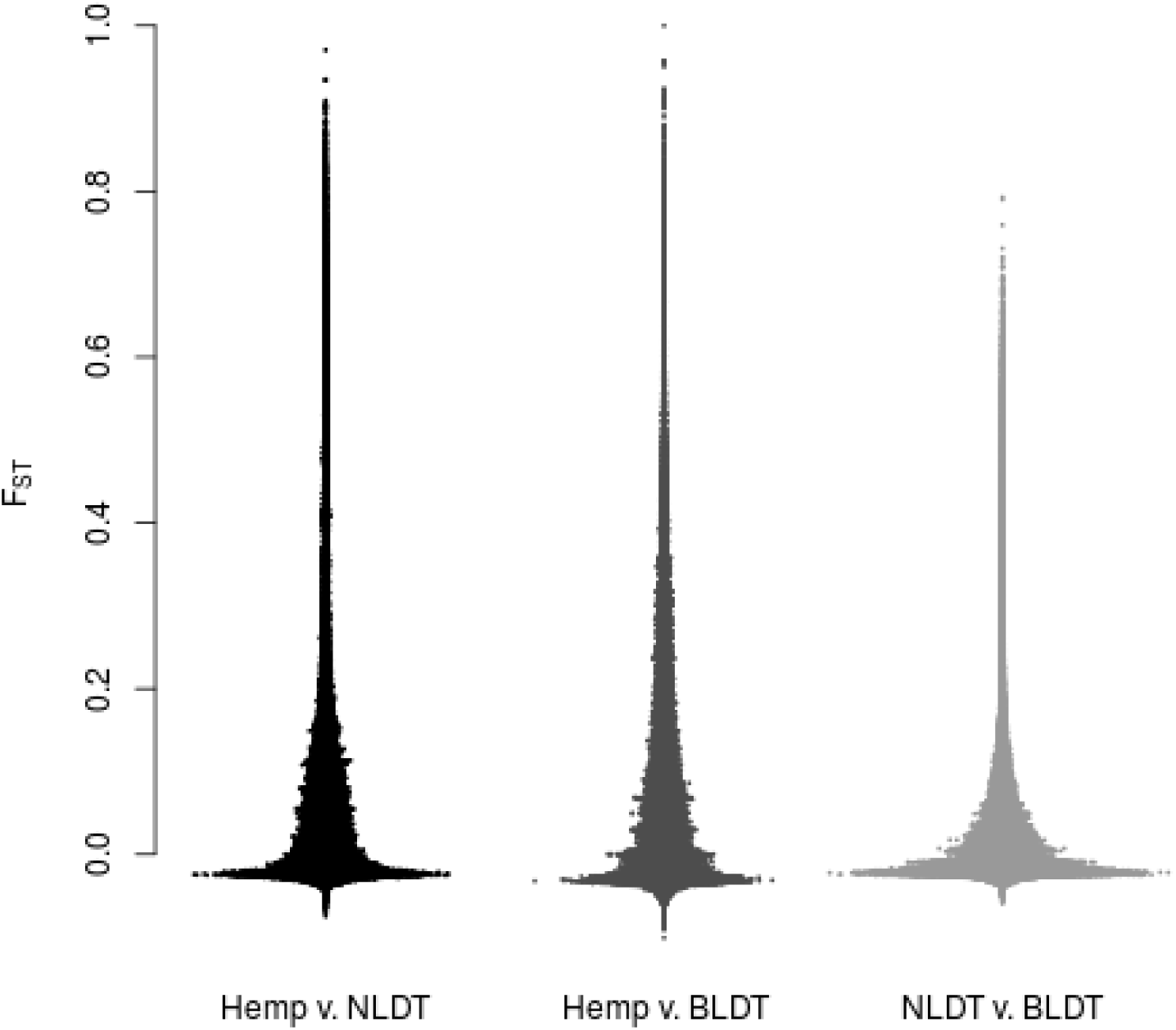
Distribution of Weir-Cockerham F_ST_ estimates for each population comparison. Each population pair has some portion of segregated sites.

Additional *Cannabis* diversity likely remains to be sampled. Notably absent from all genome sequence datasets published to date are putative *C. ruderalis* (59) samples. These are short weedy plants, with free shattering inflorescences found widely from northern Siberia, through central Asia and into Eastern Europe (60). Whether these populations represent ancestral, pre-domesticated wild *Cannabis*, more recent feral escapes or some combination of both remains unclear. Even though we were not able to sample putative *C. ruderalis* populations, Finola is an early maturing seed hemp strain from Finland with purported northern Russian landrace ancestry (52), and Low Ryder and Auto AK-47 are auto-flowering drug-type strains with possible *C. ruderalis* heritage included in our samples (Table 2). Our analyses found Finola fits within the hemp group while Low Ryder and Auto AK-47 are close relatives of each other within the NLDT group (Figure S3). Further genomic analyses are required to determine the extent to which *C. ruderalis* populations are genetically distinct from hemp and drug-type groups, and whether they may in fact harbor an ancestral wild-type gene pool from which European hemp varieties were domesticated (5, 16).

Broad leaflet Asian hemp is also underrepresented, although we included one putative Chinese hemp sample that occupies an area between the core hemp and BLDT populations (Table 2, Figure 2 and Figure S2). Hillig’s (5) analysis of alloenzymes concluded that Asian hemp strains were more similar to Asian drug-type strains than they were to narrow leaflet European hemp. Likewise, Gao et al. (61) found genetic dissimilarity between European hemp and Chinese hemp, using microsatellites, and showed at least several distinct groups of hemp occur across the vast geography of Asia. Overall, Asian and European hemp strains appear dissimilar genetically, possibly reflecting independent domestication events (60).

One major complication obscuring the understanding of *Cannabis* diversity and history is the lack of information about the native range or ranges of *Cannabis*. In addition to divergent breeding efforts and human-vectored transport of seeds, the tendency of *Cannabis* is to escape into feral populations wherever human cultivation occurs in temperate climates (62). This, coupled with wind pollination biology and no known reproductive barriers, makes the existence of pure wild native *Cannabis* populations unlikely. The weedy tendencies of *Cannabis* are exemplified by the mid-western USA populations of feral hemp that flourish despite the eradication efforts by the Drug Enforcement Agency, which have for decades totaled millions of plants removed per year. A comprehensive evaluation of *Cannabis* diversity, which includes feral and wild Eurasian populations, is required to ascertain if the levels of divergence and gene flow are consistent with one or more origins of domestication (5). Even if these extant populations are highly admixed with modern varieties, their study promises to offer insight into *Cannabis* ecology and evolution, given how different the selective regime of the feral setting is compared to that of agricultural fields. Considering the similar debates regarding the timing and origins of *Oryza* domestication that remain as of yet unresolved (63), *Cannabis* requires substantially more work to unravel its complicated relationship with humans.

‘Indica’ and ‘sativa’ are commonly used terms ascribed to plants that have certain characteristics, often related to leaflet morphology and the perceived effects of consuming the plant (8). However these names are rooted in taxonomic traditions dating to Linnaeus who first classified the genus as monotypic *(Cannabis sativa)* based on hemp specimens from Virginia and Europe (64). Lamarck subsequently designated *Cannabis indica* to accommodate the shorter stature potent narrow leaflet drug-type plants from the Indian subcontinent (65). Although currently the term ‘indica’ is typically used to refer to BLDTs, this biotype from the Hindu-Kush mountains (14) was not clearly documented until a 1929 survey of Afghani agriculture by Vavilov (66). This absence of historical documentation until the 20^th^ century, a very narrow geographic range, and some evidence for a broader NLDT gene pool (Table 1, Supplementary Figures 3 and 4), suggest a separate and more recent origin of the BLDT clade. This origin could represent a domestication event of a wild or feral BLDT population, or perhaps hybridization events between NLDT and BLDT populations. Final resolution of *Cannabis* taxonomy will require complete assessment of standing global genetic diversity and experimental evaluation of reproductive compatibility across all major genetic groups (67), in conjunction with morphological circumscriptions. Given the current absence of evidence for reproductive barriers, and overall limited genetic distances between hemp and drug-type strains analyzed in this study we suggest continued monotypic treatment of plants in this genus as *Cannabis sativa* L. is warranted.

**Cannabinoid and Terpenoid Diversity**. THCA and CBDA are the most abundant cannabinoids produced by the majority of strains on the North American market today (Figure 5a), and both compounds show an impressive range of medicinal potential (33, 68), although endocannabinoid-based therapy trials have a history of significant rates of study withdraws and adverse effects (69). Historical breeding efforts have resulted in mostly high THCA plants that produce strong intoxicating effects when consumed, and that synthesize only very low levels of alternative cannabinoids (Figure 5b). High CBDA plants have only recently become more available in North America over the last several years in response to demand. Interestingly, these high CBDA-producing plants form several clusters within the both the NLDT and BLDT groups, as well as within the hemp group (Dataset S1), but rarely reach equivalent quantities total cannabinoid production as those found in high THCA plants (Figure 4a). The minor cannabinoids that are commonly assayed, CBGA, CBCA, THCVA and CBDVA are also of interest, despite strains producing high levels of these compounds being largely unavailable for research currently (70). With at least 74 cannabionoids identified in *Cannabis*, modernized genetic and breeding techniques are required to diversify and optimize *Cannabis* varieties. Efforts should also be made to document and preserve feral, wild and heirloom populations that can serve as reservoirs of cultural and genetic diversity.

Aromatic terpenoids impart many of the characteristic fragrances to *Cannabis*, and possibly contribute to the effects of consumption (2). Terpenoids are synthesized in many plant species, and play a role in relieving various abiotic and biotic stresses through direct and indirect mechanisms (71). Our analysis of strains sharing common genetic groups shows that each group has a distinct terpenoid profile (Figure 5c and Figure S5). We found NLDTs to contain significantly more β-myrcene and α-terpinolene than BLDTs, although interestingly the two hemp strains for which we analyzed chemical data for had significantly more β-myrcene than either drug-type group (Figure 5c). Similarly Hillig (72) found NLDTs to yield significantly more β-myrcene than Afghani BLDTs, yet European hemp and un-cultivated accessions labeled as *C. ruderalis* contained the highest levels. Hillig also reported that Afghani BLDTs contained the highest levels of guaiol and eudesmol isomers, which we did not measure, although we found BLDTs contained more linalool than NLDTs or hemp. Understanding the ecological functions and evolutionary origins of terpenoids and cannabinoids in *Cannabis* could improve therapeutic potential, and possibly reduce the need for pesticide application during cultivation.

**Figure 5.**
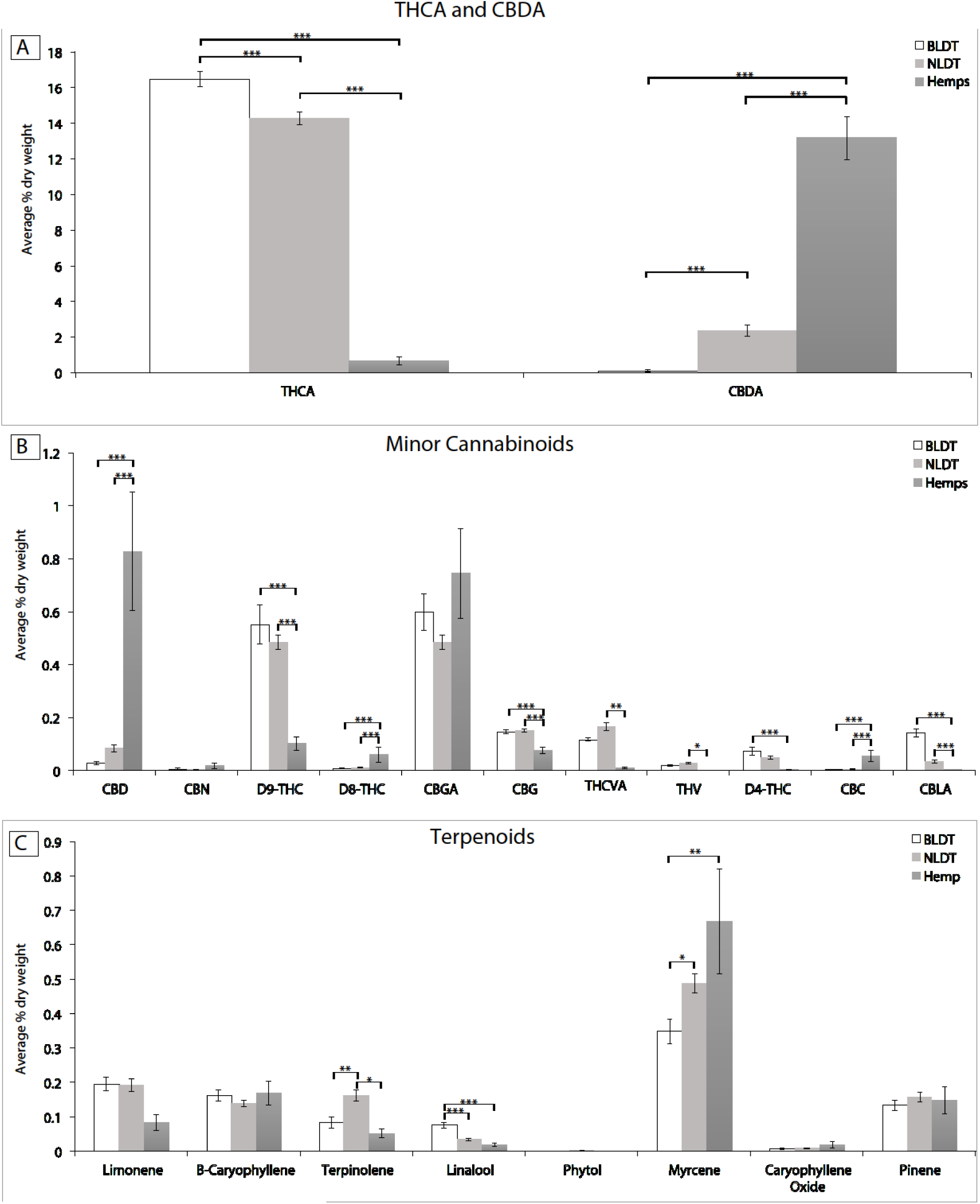
Average percentage of mass for dried and un-pollenated female flowers of *Cannabis* genetic groups. (a) THCA and CBDA cannabinoids (b) Minor cannabinoids (c) Terpenoids. THCA = delta-9-tetrahydrocannabinolic acid. CBDA = cannabidiolic acid. CBD = cannabidiol. CBN = cannabinol. D9-THC = delta-9-tetrahydrocannabinol. D8-THC = delta-8-tetrahydrocannabinol. CBGA = cannabigerolic acid. CBG = cannabigerol. THCVA = Tetrahydrocannabivarin carboxylic acid. THCV = Tetrahydrocannabivarin. D4-THC = delta-4-tetrahydrocannabinol. CBC = cannabichromene. CBLA = cannabicyclolic acid.

**Conclusions**. *Cannabis* genomics offers a window into the past, but also a road forward. Although historical and clandestine breeding efforts have been clearly successful in many regards (21, 31), *Cannabis* lags decades behind other major crop species in many other respects. Developing stable *Cannabis* lines capable of producing the full range of potentially therapeutic cannabinoids is important for the research and medical communities, which currently lack access to diverse high-quality material in the USA (73).

In this paper we extended the initial *Cannabis* genome study (10), by re-mapping WGS and GBS sequence reads to the existing PK draft scaffolds, to understand diversity and evolutionary relationships among the major lineages. Although hybridization of cultivated varieties (52) and human transport of seeds across the globe was hypothesized to have obscured much of the ancestral genetic signal (13), we found significant evidence for apparent ancestral signals in genomic data derived largely from modern cultivated varieties (Table 2, Figures 2 and 3). Re-analysis of previously published GBS data (50) provides additional limited evidence for a fourth group (Supplementary Figures 4 and 5). Interestingly, unique cannabinoid and terpenoid profiles were associated with three of the genetic groups, lending support to their validity, despite the limitations of our sampling scheme. Overall, we hope the publicly available data and analyses from this study will facilitate the continued research on the history of this controversial plant and the development of the agricultural and therapeutic potential of *Cannabis*.

## Materials and Methods

**Sample collection**. DNA was obtained from numerous sources, including a variety of breeding and production facilities. The strain names, descriptions and putative origins used in this paper were recorded from the providers of the DNA and sequence data (Dataset S1). For data not previously published, DNA extractions were performed using the Qiagen DNeasy Plant Mini Kit (Valencia, CA) according to the manufacturer’s protocol.

**Whole genome shotgun (WGS) sequencing**. 60 samples were sequenced using standard Illumina multiplexed library preparation protocols for two 2 x 125 HiSeq 2500 lanes and one 2 x 150 NextSeq 500 run. Sequencing efforts were targeted for approximately 4-6x coverage of the *Cannabis* genome per sample.

**Genotype-by-Sequencing (GBS)**. 182 samples were sequenced on two 1 x 100 HiSeq 2500 lanes, following a multiplexed library preparation protocol described previously (74).

**Publically available data**. We obtained three WGS datasets available from NCBI (10) and received seven additional WGS datasets from Medicinal Genomics Corporation (www.medicinalgenomics.com). GBS data for 143 samples from Sawler et al. (50) were also included in this study.

**Sequence Processing, Alignment and SNP calling**. Trimmomatic (75) was used to trim any remaining adaptor sequence from raw fastq reads and remove sequences with low quality regions or ambiguous base calls using the following settings:

ILLUMINACLIP:IlluminaAdapters:2:20:10 LEADING:20 TRAILING:20

SLIDINGWINDOW:5:15 MINLEN:100. Trimmed raw reads from the 67 total WGS samples were then aligned to the only publicly available draft genome of PK (JH226140-JH286168) using the Burrows-Wheeler Alignment tool (BWA mem) (76). Chloroplast and mitochondrial regions were excluded. We collated the individual alignments to produce a single variant call format table (.vcf) for all samples using samtools mpileup -uf | bcftools view -bvcg (77). We filtered the vcf table to include only high quality informative SNP sites using vcftools (78), bash and awk with the following vcf parameters: Q ( >200), GQ (>10), AF1 (.1 - .9), biallelic sites only and no ambiguous bases. Next, data filters were applied through plink (79) to require that individuals have a minimum 50% informative sites and that sites each have data for minimum 20% of samples. Finally we used an estimate of expected coverage for the single copy portion of the genome based on the estimated genome size and number of reads being aligned. This was adjusted empirically based on total coverage level (across all WGS samples) per SNP site (Figure S1) and bounded by a 95% Poisson confidence interval (mean 362x coverage). Further removal of repetitive content was achieved by aligning the PK reference to itself with BLASTN and removing all sites that were within regions of ≥ 97% identity for at ≥ 500 bp alignments. These aforementioned processing, alignment and SNP calling procedures were then preformed separately on the 182 GBS samples generated for this study and the 143 GBS samples previously published (50), resulting in three vcf tables and filtered SNP sets. GBS SNPs were additionally required to have a minimum of 5x coverage per sample. Due to limited overlap between the SNP sites produced by the two GBS libraries, most downstream analyses were performed separately for each GBS library along with its corresponding set of WGS SNPs. Code used for these analyses is available at https://github.com/KaneLab.

**SNP Analyses**. To visualize genetic relationships, divergence, and ancestral hybridization among lineages, a phylogenetic neighbor network was inferred using simple p-distance calculations (51). Heterozygosity counts and Multidimensional Scaling (MDS) analyses were calculated with Plink (79). Average within and between group genetic distances, and a 45 SNP alignment neighbor joining tree based on p-distances, were calculated with MEGA6 (80). Population structure inferences were made through FastStructure (54) and FLOCK (53). Tests for reticulation within the trees and admixture between populations were performed in TreeMix (81) F_ST_ estimates were calculated with vcftools (78).

**Chemical Analyses of Genetic Groups**. The cannabinoid and terpenoid information (chemotype) for a portion of the strains in the genome analysis were generated by Steep Hill Labs (http://steephill.com/). Only strains with matching data in the genomic analysis were analyzed, for a total of 112 individuals from 17 strains from the BLDT group, 278 individuals from 35 unique strains from the NLDT group, and 33 individuals from two strains of hemp, for a total of 423 individuals in this analysis (Dataset S1). This chemotype analysis was performed using high performance liquid chromatography (HPLC) with Agilent (1260 Infinity, Santa Clara, CA) and Shimadzu (Prominence HPLC, Columbia, MD) equipment. Between 400 and 600 milligrams of each sample was extracted into methanol, diluted and analyzed by HPLC. A mobile phase consisting of 0.1% formic acid in water and 0.1% formic acid in methanol was used with a gradient starting at 72% methanol and ending at 99% methanol. Terpenoid standards were purchased from Sigma-Aldrich (St. Louis, MO). Cannabinoid standards were purchased from Cerilliant (Round Rock, TX), RESTEK (Bellefonte, PA) and Lipomed (Cambridge, MA).

A C18 column from RESTEK (Raptor ARC-18, Bellefonte, PA) or Phenomenex (Kinetex C18, Torrance, CA) was used. Concentrations of cannabinoids without commercially available standards were estimated using published absorptivies (82). The chemotype data analyzed for this research includes 13 cannabinoids and eight terpenoids. Each compound was quantified using a linear calibration curve. Analytes were measured as percent mass in sample and not corrected for moisture content.

We performed a one-way ANOVA for each cannabinoid and terpenoid separately, with the group (NLDT, BLDT, and hemp) as the predictor variable. We used Bonferroni corrections for multiple comparisons. We also implemented a Principal Component Analysis (PCA) with prcomp function in base R, and car was used to visualize 95% confidence ellipses for each group (www.R-project.org). Individuals with missing data values for any cannabinoid or terpenoid were removed. After removing the individuals with missing values, we had a total of 351 individuals: 94 BLDT, 229 NLDT, and 28 hemp.

## Acknowledgments.

We thank Ben Holmes of Centennial Seeds; Devin Liles, Carter Casad and Jan Cole of The Farm; Ashley Edwards of Ward, Colorado; Jake Salazar of MMJ America; Kevin McKernan of Medicinal Genomics; David Salama, Ashley and Matt Rheingold of Headquarters; Ezra Huscher; Nico Escondido and Bob Sievers for providing DNA samples. We thank Reggie Gaudino of Steep Hill for advice and assistance with the chemical data. This project was supported by donations to the University of Colorado Foundation gift fund 13401977-Fin8 to NCK.

Author contributions. RCL, DV, KHW and NCK designed the project. CJS and MJG collected samples. KHW generated DNA sequencing libraries. RCL, NCK and SBT performed bioinformatics analyses. KdC, DPL and TCR generated chemical data. DV and SBT performed chemical data analyses. RCL, DV and NCK wrote the paper.

## Supplementary Information

**Figure S1.**
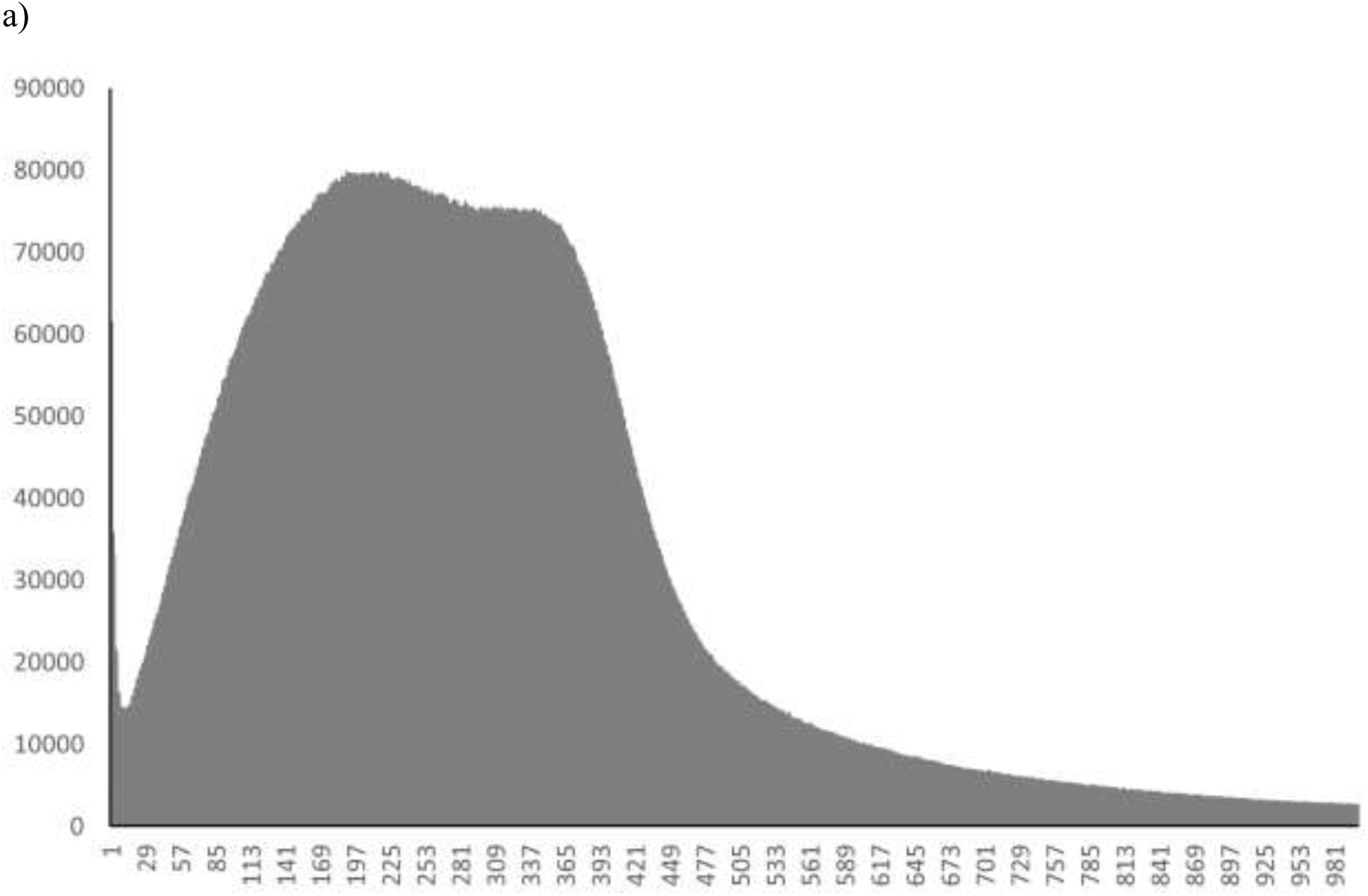
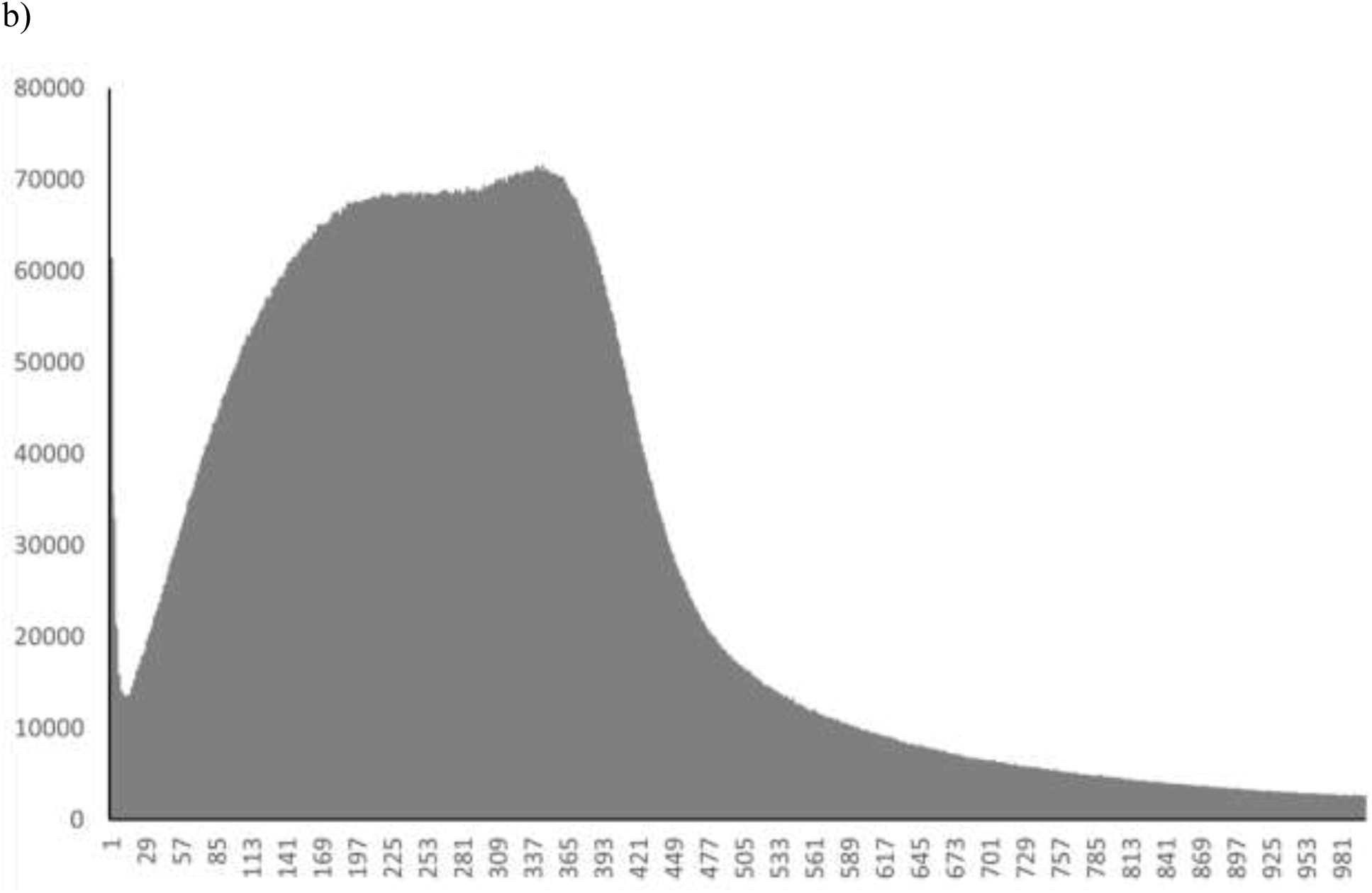
Histograms of WGS read depths at variant loci. a) before PK reference self-similarity filter. b) after self-similarity filter.

**Figure S2.**
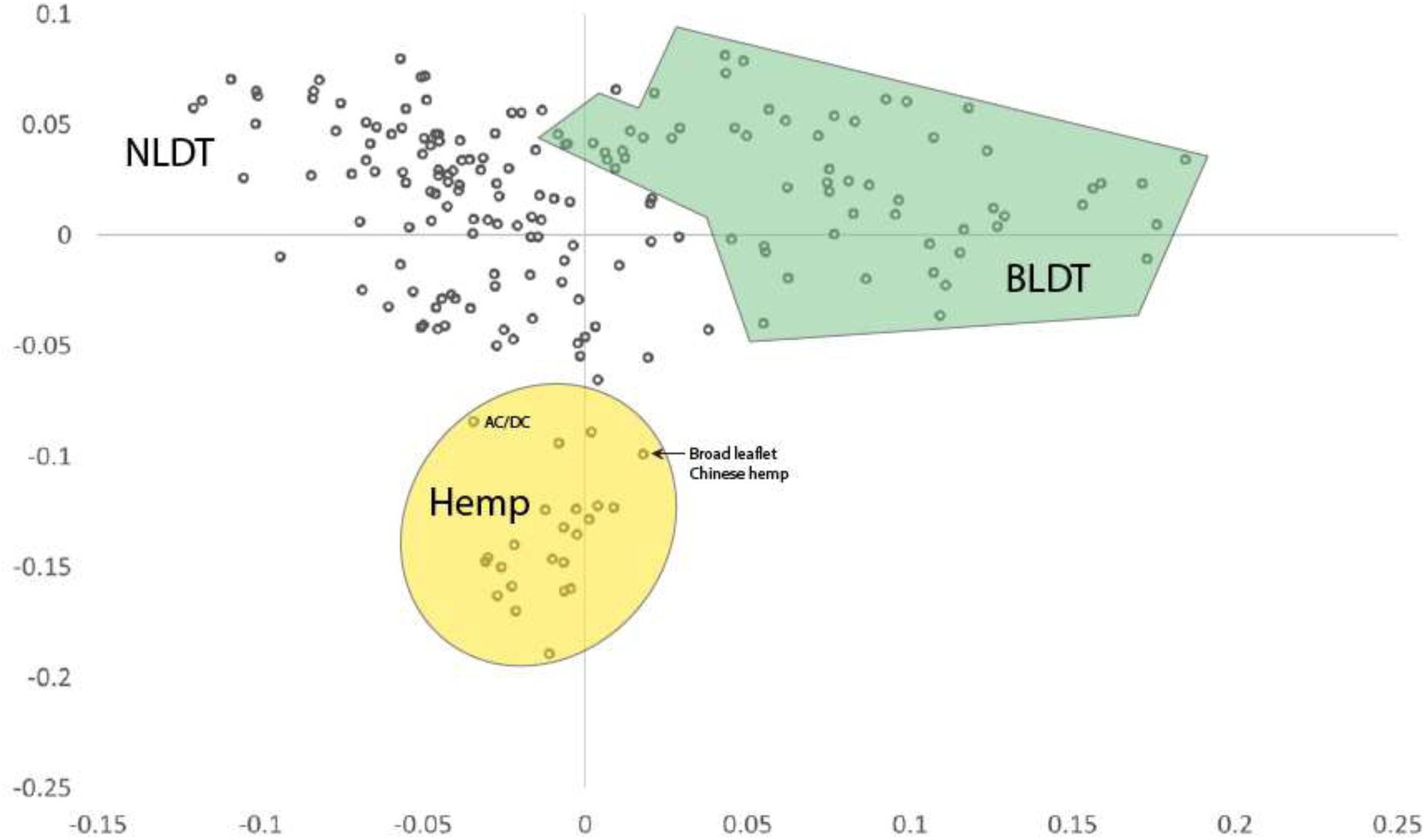
Multidimensional Scaling plot of GBS and WGS SNPs. Hemp, NLDT and BLDT group assignments were made by FLOCK.

**Figure S3.** Phylogenetic neighbor network of WGS samples combined with Sawler GBS SNPs (4,105) in separate high resolution pdf.

**Figure S4.** Neighbor joining tree from 45 SNP alignment of 289 GBS and WGS samples in separate high resolution pdf.

**Figure S5.**
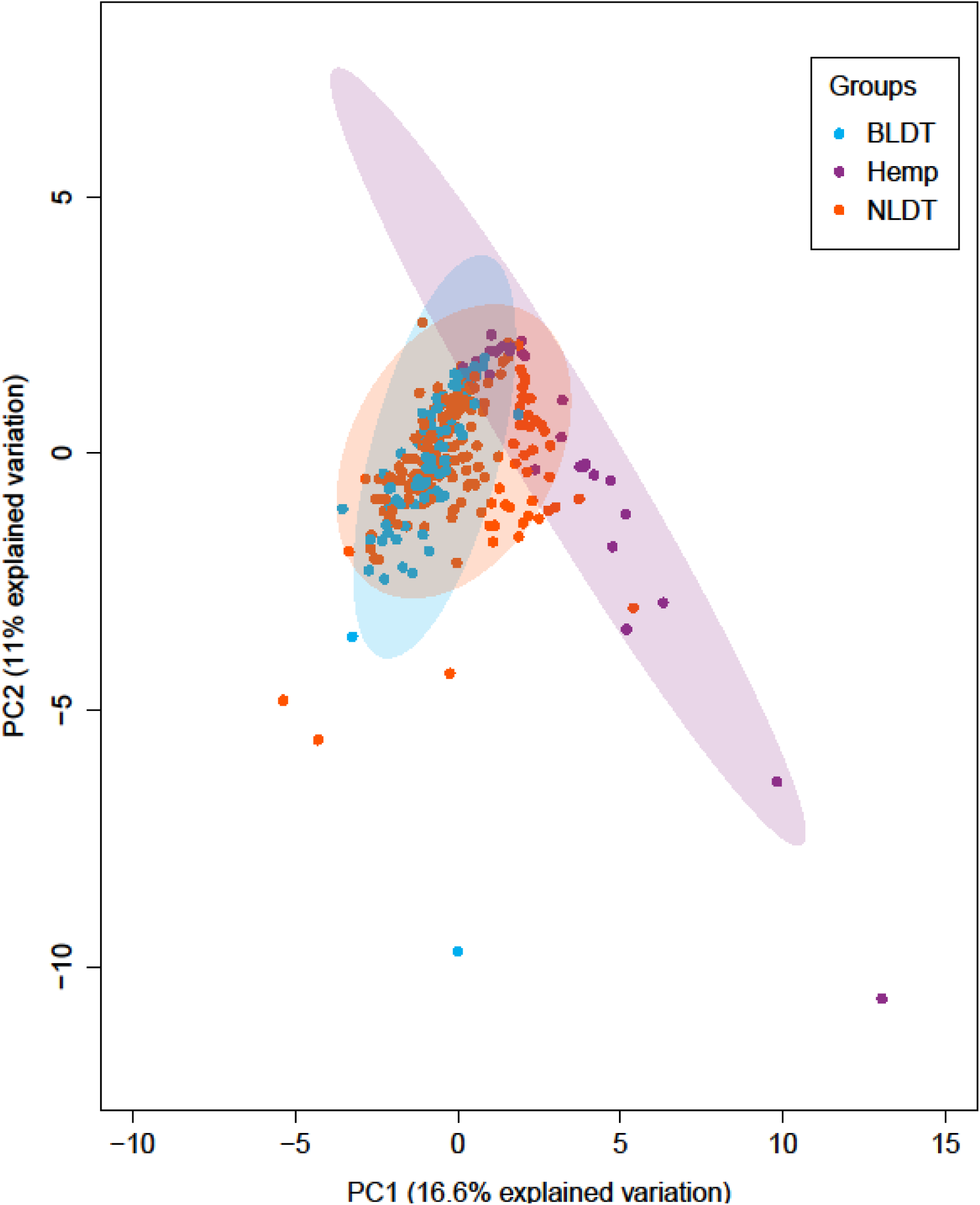
Principal Components Analysis of cannabinoid and terpene profiles colored by FLOCK derived genetic groups.

